# A comparative genomics approach reveals a local genetic signature of *Leishmania tropica* in Morocco

**DOI:** 10.1101/2023.05.25.542267

**Authors:** Hasnaa Talimi, Othmane Daoui, Giovanni Bussotti, Idris Mhaidi, Anne Boland, Jean-François Deleuze, Rachida Fissoune, Gerald F. Späth, Meryem Lemrani

## Abstract

In Morocco, cutaneous leishmaniasis (CL) caused by *Leishmania* (*L.*) *tropica* is an important health problem. Despite its high incidence in the country, the genomic heterogeneity of these parasites is still incompletely understood. In this study, we sequenced the genomes of 14 Moroccan isolates of *L. tropica* collected from confirmed cases of CL to investigate their genomic heterogeneity. Comparative genomics analyses were conducted by applying the recently established Genome Instability Pipeline (GIP), which allowed us to conduct phylogenomic and PCA analyses, and to assess genomic variations at the levels of the karyotype, gene copy number, and single nucleotide polymorphisms (SNPs). The results identified a core group of 12 isolates that were genetically highly related but evolutionarily distant to the reference genome as judged by the presence of over 100,000 SNPs, 75% of which were shared inside this core group. In addition, we identified two highly divergent strains, M3015 and Ltr_16, that were phylogenetically distinct between each other as well as to the core group and the reference genome. Read-depth analysis revealed important karyotypic variations across all isolates and uncovered important differences in gene copy number between the isolates of the core group and the *L. tropica* reference genome, as well as between the core group and M3015. In conclusion, our NGS results suggest the presence of a local SNP signature that distinguishes Moroccan *L. tropica* from other endemic regions and from the reference genome. These results pave the way for future research with a larger number of strains that will allow to correlate diverse phenotypes (resistance to treatments, virulence) and origins (geography, host species, year of isolation) to defined genomic signals that may represent interesting biomarker candidates.

## INTRODUCTION

Leishmaniasis is a vector-borne disease of global reach with important public health impact. The disease is of zoonotic or anthroponotic origin and is caused by protozoan parasites of the genus *Leishmania* that show cutaneous or visceral tropisms. The parasite is transmitted by the bites of infected female phlebotomine sandflies, which feed on blood [1]. Cutaneous leishmaniasis (CL) is the most frequent form of the disease, associated with skin lesions, primarily ulcers, on exposed regions of the body, resulting in life-long scars that can cause social stigma [2]. The Americas, the Mediterranean Basin, the Middle East, and Central Asia account for around 95% of all CL cases. In 2020, ten countries accounted for more than 85% of new CL cases: Afghanistan, Algeria, Brazil, Colombia, Iraq, Libya, Pakistan, Peru, the Syrian Arab Republic, and Tunisia. Every year, between 600,000 and 1 million new CL cases are reported globally [3]. The epidemiology of leishmaniasis is determined by both parasite and sandfly species, local ecological factors of transmission locations such as animal reservoir or climatic conditions, present and previous parasite exposure of the human population, and human behavior [3]. Approximately 70 mammal species, including humans, have been identified as natural reservoir hosts of *Leishmania* parasites [3].

In Morocco, CL is caused by three species of *Leishmania*: *L. major, L. infantum* and *L. tropica*. The latter is the main causative agent of CL due to its high frequency and wide geographical distribution. CL caused by *L. tropica* was described for the first time in 1987 in a child who had resided in the province of Azilal, central Morocco [4]. Thereafter, a large hypo-endemic focus between Tadla and Agadir has been identified [5]. Since the mid of 1990s, CL outbreaks due to *L. tropica* occurred in emerging foci in central and northern Morocco, and *L. tropica* was even found in areas previously only known as *L. major* foci [6–8]. In Morocco, the large genetic heterogeneity of *L. tropica* is one of its most distinctive features, as judged by isoenzymatic variability, with multilocus enzyme electrophoresis (MLEE) distinguishing ten distinct zymodemes [5, 9]. This polymorphism was also revealed by random amplification of polymorphic DNA (RAPD), PCR-RFLP of the antigen-encoding genes gp63 and cpb, sequence analysis of intergenic spacer regions (ITS), multilocus microsatellite typing (MLMT), and multilocus sequence typing (MLST) [10, 11].

Next Generation sequencing (NGS) analysis applied on *Leishmania* clinical isolates changed the way we approach problems in basic and epidemiological research. The ability to sequence the entire genome of many related organisms has enabled large-scale comparative and evolutionary studies [12] and for example allowed to uncover mechanisms of *Leishmania* genomic adaptation, thus providing novel insight into genome complexity and gene dosage-dependent gene regulation [13–17]. NGS also has been successfully applied in drug resistance research in *Leishmania*, which correlated changes in chromosome copy number, gene dosage, and SNP frequencies to resistance in both laboratory-and field-derived strains of *Leishmania* [18, 19].

Here, we performed a comparative genomics analysis on 14 novel genomes of *L. tropica* that were isolated from confirmed cases of CL in Morocco. We used the Genome Instability Pipeline (GIP) [20] to assess the diversity between isolates and with respect to the reference genome, and to evaluate the genetic heterogeneity between isolates by analyzing the differences in karyotype, gene copy number, and single nucleotide polymorphisms (SNPs).

## METHODS

### Leishmania isolates and parasite culture

The ethics protocols of sampling differ depending on whether one is working with humans or dogs. A sampling of human patients involves obtaining informed consent and following ethical guidelines in accordance with the principles of medical ethics, e.g., using sterile techniques. While the sampling of dogs may involve some ethical considerations, it is not as complex as that for human patients which is obtained by non-invasive methods, such as scraping or swabbing the skin lesions. Within the context of the present study, all adult participants provided informed consent prior to their involvement. For the inclusion of young children, consent was obtained from their respective parents or legal guardians. The study underwent thorough evaluation and received approval from the Ethical Review Committee for Biomedical Research (CERB) of the Faculty of Medicine and Pharmacy, Rabat, Morocco (IORG 0006594 FWA 00024287), in adherence to established research guidelines.

We included in this study 14 isolates of *L. tropica* from confirmed cases of CL in Morocco, which were collected from 13 patients and one dog between 1990 and 2020 in various regions of Morocco, including Essaouira, Ouarzazate, and the macrofocus of the Azilal province (Foum-Jemaa, Azilal, and Tanant) (Table 1 and Figure 1). Only one isolate showed relapse.

**Figure 1.**
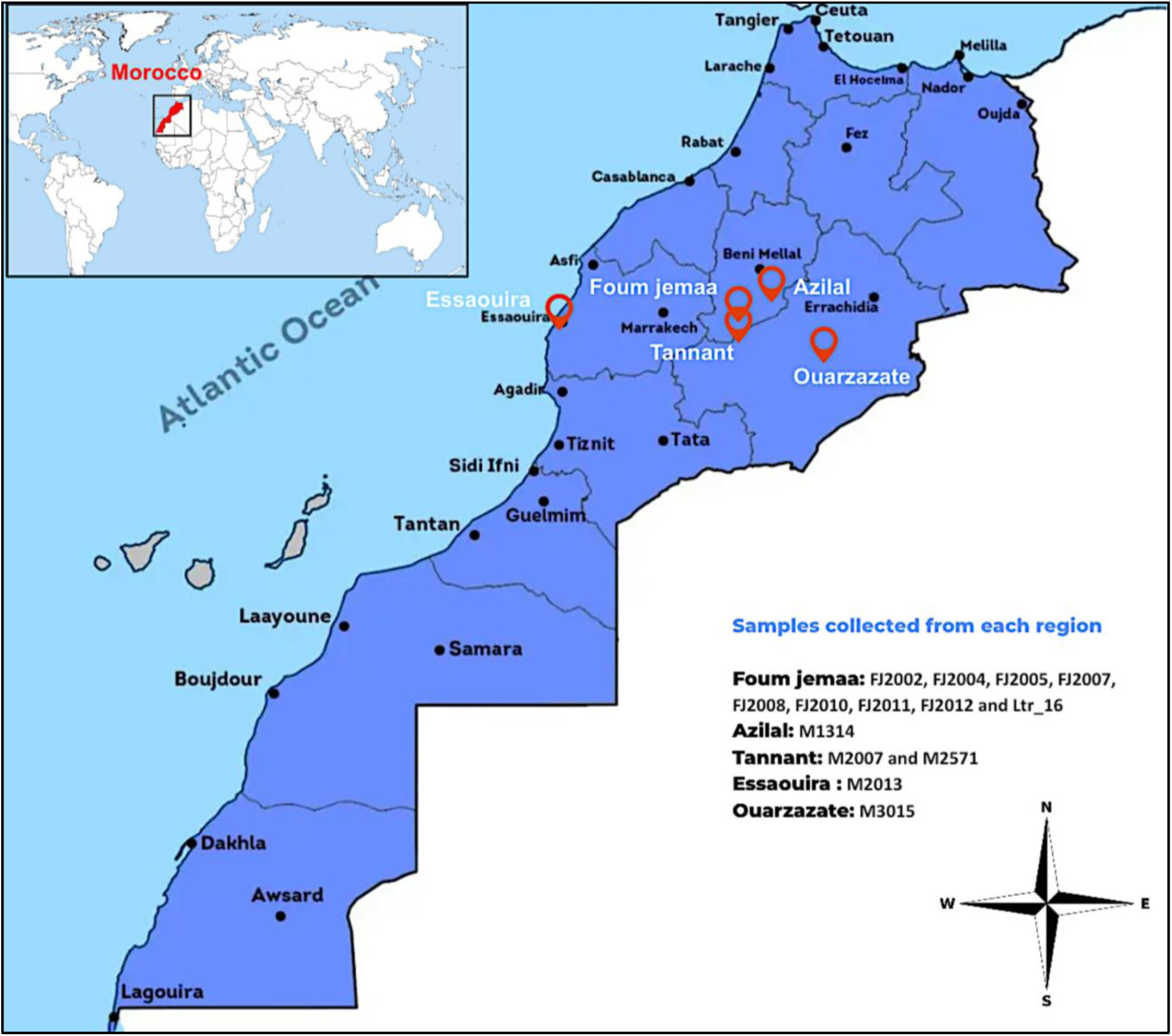
Map showing the geographic origin of the isolates studied.

**Table 1.** Overview of isolates.

| **Sample ID** | **Host / Pathology** | **Origin** | **Year of isolation** | **Specimen** | **Zymodeme** | **Molecular typing** | **Passage** | **Circumstances of sampling** |
| --- | --- | --- | --- | --- | --- | --- | --- | --- |
| **FJ2002** | Human / CL | Foum jemaa | 2020 | skin tissue | n.d. | ITS1-PCR-RFLP | < 5 | Cured |
| **FJ2004** | Human / CL | Foum jemaa | 2020 | skin tissue | n.d. | ITS1-PCR-RFLP | < 5 | Cured |
| **FJ2005** | Human / CL | Foum jemaa | 2020 | skin tissue | n.d. | ITS1-PCR-RFLP | < 5 | n.d. |
| **FJ2007** | Human / CL | Foum jemaa | 2020 | skin tissue | n.d. | ITS1-PCR-RFLP | < 5 | Relapse |
| **FJ2008** | Human / CL | Foum jemaa | 2020 | skin tissue | n.d. | ITS1-PCR-RFLP | < 5 | Cured |
| **FJ2010** | Human / CL | Foum jemaa | 2020 | skin tissue | n.d. | ITS1-PCR-RFLP | < 5 | Cured |
| **FJ2011** | Human / CL | Foum jemaa | 2020 | skin tissue | n.d. | ITS1-PCR-RFLP | < 5 | n.d. |
| **FJ2012** | Human / CL | Foum jemaa | 2020 | skin tissue | n.d. | ITS1-PCR-RFLP | < 5 | Cured |
| **Ltr_16** | Human / CL | Foum jemaa | 2016 | skin tissue | n.d. | ITS1-PCR-RFLP | n.d. | Cured |
| **M1314** | Human / CL | Azilal | 1988 | skin tissue | MON - 109 | ITS1-PCR-RFLP | n.d. | n.d. |
| **M2007** | Canine/ CanL | Tannant | 1990 | skin tissue | MON - 102 | ITS1-PCR-RFLP | n.d. | n.d. |
| **M2013** | Human / CL | Essaouira | 1990 | skin tissue | MON - 102 | ITS1-PCR-RFLP | n.d. | n.d. |
| **M3015** | Human / CL | Ouarzazate | 1995 | skin tissue | MON - 264 | ITS1-PCR-RFLP | n.d. | n.d. |
| **M2571** | Human / CL | Tannant | 1993 | skin tissue | MON - 102 | ITS1-PCR-RFLP | n.d. | n.d. |

Sampling of isolates with identifiers beginning with “FJ” was carried out by dermal scraping of the lesion edge, whereas the ones with identifiers starting with “M” were received from the Parasitology, Mycology Department, CHU de Montpellier (Site Antonin Balmes-La Colombière). Parasite isolates were first grown in biphasic NNN (Novy-MacNeal-Nicolle) medium at 25°C and established cultures were expanded for 2 to 5 passages in RPMI 1640 (Biowest, Nuaille, France) medium supplemented with 2 mM L-glutamine (Eurobio, Les Ulis, France), 15% fetal bovine serum (Biowest, Nuaille, France), and 1% penicillin/streptomycin (100 U/mL penicillin and 100 μg/mL streptomycin; Biowest, Nuaille, France).

**DNA extraction and sequencing analysis**.

Cryopreserved *Leishmania* promastigotes were thawed and first cultured at 25 °C in NNN medium before grown in RPMI 1640 medium (Biowest, Nuaille, France). DNA extraction was performed using QIAamp DNA Mini Kit (Qiagen, Germany) according to manufacturer instructions. DNA quality and quantity were assessed using NanoDrop (Thermo Fisher Scientific, Waltham, MA, USA).

Whole genome sequencing was performed by the Centre National de Recherche en Génomique Humaine (CNRGH, Institut de Biologie François Jacob, CEA, Evry, FRANCE). After quality control, genomic DNA (1µg) was used to prepare the library for whole genome sequencing on an automated platform, using the Illumina “TruSeq DNA PCR-Free Library Preparation Kit”, according to the manufacturer’s instructions. After normalization and quality control, qualified libraries were sequenced on a HiSeqX5 platform from Illumina (Illumina Inc., CA, USA), as paired-end, 150 bp reads. 24 samples have been pooled on one lane of the HiSeqX5 flow cell. Sequence quality parameters have been assessed throughout the sequencing run and standard bioinformatics analysis of sequencing data was based on the Illumina pipeline to generate a FASTQ file for each sample.

### Comparative genomic analyses

Reads were mapped against the Leishmania_tropica_CDC216-162 reference genome [21]. Genome annotation (gff3 file format) was performed with the Companion software package (http://companion.gla.ac.uk/), an online server designed for the annotation and analysis of genomes [22]. We used transcriptome evidence generated with Cufflinks and LtropicaL590 as a reference organism with default pseudo-chromosome settings. All data are available under URL: https://companion.gla.ac.uk/jobs/afa429f7f98eb14d3e4db906.

The GIP pipeline was developed by integrating various commonly used bioinformatic tools for genetic analysis. For instance, within the pipeline, Freebayes (version 1.3.2) was employed to detect single nucleotide polymorphisms (SNPs) using the following parameters: “--read-indel-limit 1 --read-mismatch-limit 3 --read-snp-limit 3 --min-alternate-fraction 0.05 --min-base-quality 5 --min-alternate-count 2 --pooled-continuous”. Additionally, snpEff (version 4.3t) was utilized to predict the potential impacts of these identified SNPs. As well as plotting the coverage ratios of the genomic bins which allows researchers to visualize the distribution of coverage across the genome and identify regions with higher or lower coverage.

All results presented in this study were generated using the Genome Instability Pipeline (GIP, [20]) version 1.0.9. The GIP code is maintained and freely distributed at the GitHub page under https://github.com/giovannibussotti/GIP. All GIP outputs were used as input for giptools, a suite of thirteen modules that allows comparing GIP-processed samples and visualizing changes in chromosomal copy number, gene copy number, and SNPs. Complete documentation for GIP and giptools, including a description of all options, is available at https://gip.readthedocs.io/en/latest/.

## RESULTS

### Assessing the genetic heterogeneity of Moroccan L. tropica field isolates reveals differences to the reference genome

We first applied cluster analyses to assess the relationship between the *Leishmania* isolates based on genomic similarity (Figure 2A and 2B). *L. tropica* (Leishmania_tropica_CDC216-162) was used as the reference strain, while *L. aethiopica and L. major* were used as outgroup genomes, which allowed us to confirm the species of our samples and to assess the evolutionary relationship to other parasite species. As judged by the monophyletic organization, all strains clustered with the *L. tropica* reference (Figure 2A), which confirmed prior species identification by isoenzyme and molecular typing. In contrast, a more detailed phylogenetic analysis between the strains and the *L. tropica* reference alone identified two strains (M3015 and Ltr_16) branching far from the reference and the other strains clustering together but independent from the reference (Figure 2B). Results clearly show a significant genomic difference between the strains compared to the reference *L. tropica*, and reveals genetically highly heterogeneous *L. tropica* strains circulating in Morocco.

**Figure 2.**
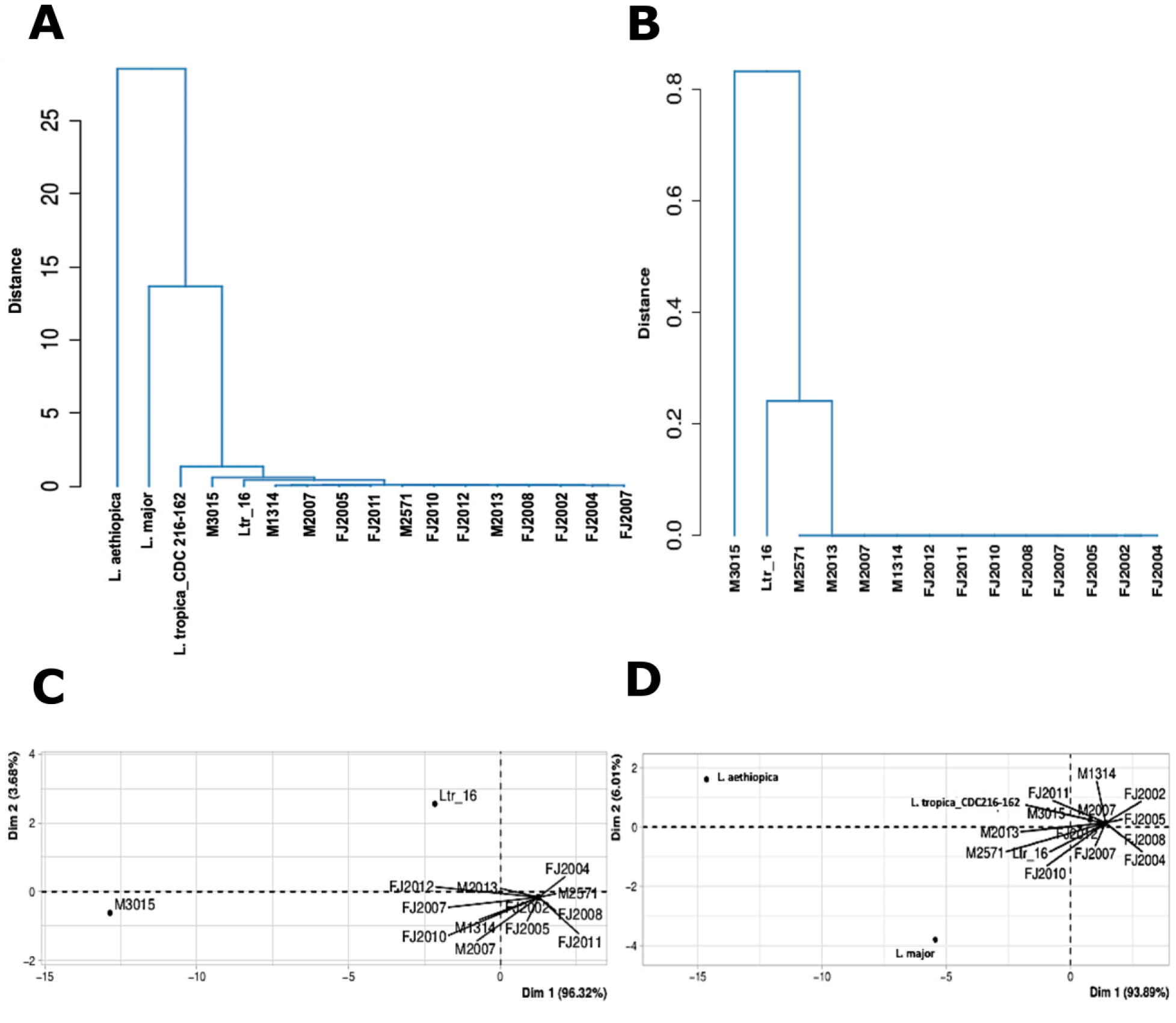
Genomic relationship between Moroccan *L. tropica* isolates. Phylogenetic tree reconstruction with added reference genomes for L. major and L. aethiopica as outgroups (A) and between the 14 isolates without outgroup (B). PCA analysis of the genetic distances estimated from the tree without (C) and with the outgroup reference genomes (D).

PCA analysis allowed us to further investigate the genomic relationship between the strains: the strains isolated in the early 1990s (M1314, M2007, M2013, and M2571) and recently in 2020 (FJ2002, FJ2004, FJ2005, FJ2007, FJ2008, FJ2010, FJ2011, and FJ2012) were grouped together, while two strains - M3015 (with rare zymodeme MON- 264, see Table 1) isolated in 1995 and Ltr_16 isolated in 2016 - were clustered separately (Figure 2C and 2D). Given the limited number of isolates, no statistically significant correlation between geography, year of isolation and genetic distance can be drawn.

### Moroccan L. tropica isolates show karyotypic variations

A comparison of chromosome numbers (somy scores) between isolates revealed significant karyotypic differences between the isolates (Figure 3A). As previously observed in all *Leishmania* species [16, 18, 23, 24], chromosome 31 was tetrasomic in all isolates, except for FJ2011 showing pantasomie. Chromosomes 33, 9, 25, 20, 8, 21, 1, and 30 were present in trisomic states in FJ2005, FJ2007, FJ2010, FJ2011, M2013, and M3015 (Figure 3A). The other chromosomes showed values between 2 and 2.5, indicating a largely disomic state for most of the chromosomes in these isolates, and revealing the presence of mosaic aneuploidies, which are likely caused by the co-existence of cell sub-populations with different somy scores as previously described in *L. donovani* [16].

**Figure 3.**
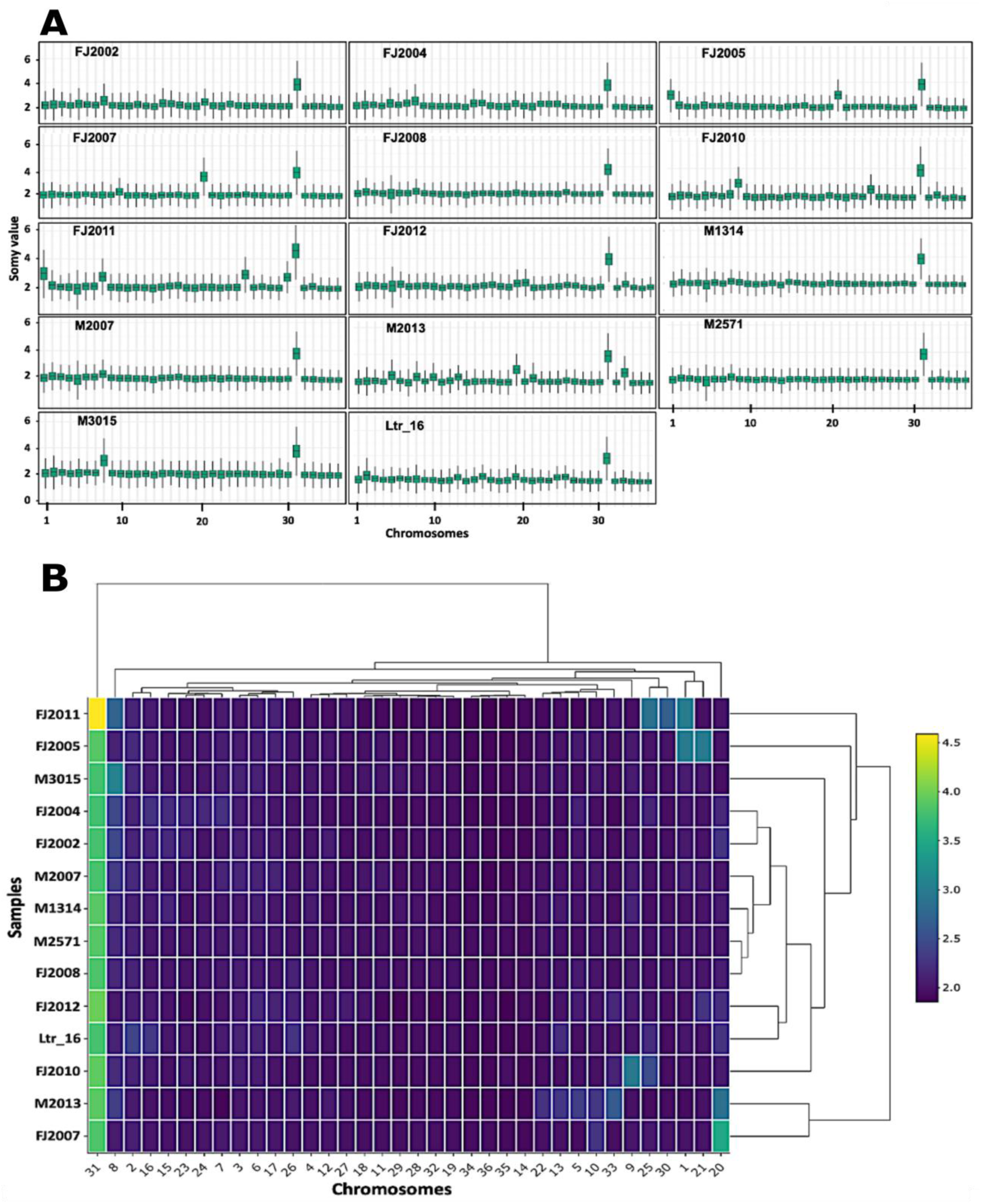
Assessment of the karyotypic profile. (A) The box plots represent the somy score distributions for each chromosome in the indicated samples. The boxes are colored according to the samples. (B) The heatmap shows the estimated chromosomal somy score for each chromosome and isolate. The color corresponds to the somy score as indicated in the legend. The dendrogram identifies distinct clusters based on aneuploidy pattern and was constructed using Euclidean distances.

The dendrogram in Figure 3B shows the results of hierarchical clustering of isolates based on their chromosomal somy patterns, which led to the identification of four distinct groups of isolates. The first group comprises M2013 (1990) and FJ2007 (2020), the second group consists of FJ2012 (2020) and Ltr_16 (2016), the third group includes FJ2004 (2020) and FJ2002 (2020), and the fourth group comprises M2007 (1990), M1314 (1988), M2571 (1993), and FJ2008 (2020). The remaining isolates (FJ2005, FJ2010, FJ2011, and M3015) are located outside the dendrogram due to their different karyotypic profile compared to the other isolates (Figure 3B). Together these data demonstrate a level of genome instability in *L. tropica* that is similar to other *Leishmania* species [13, 24]. Whether these karyotypic changes were selected in the field or are the result of culture adaptation remains to be established.

### Analysis of copy number variation

In order to further evaluate the genetic relationship between the core group and the outlier, a comparison of the sequencing coverage of FJ2005 and M3015 isolates was conducted by plotting the coverage ratio of normalized genomic bins. Specifically, we selected FJ2005 as a representative isolate for the core group and M3015 as the highly distinct isolate (Figure 4A). Comparing read depth ratios between FJ2005 and M3015 revealed 1292 and 1031 bins that were either amplified or depleted between the isolates. The ratio scores for each genomic position are shown in Supplementary Tables S2 and S3. Analyzing gene CNVs between these two isolates revealed three distinct groups of genes: (i) those whose copy number are centered around a ratio of 1, indicating a similar copy number as in the reference genome, (ii) genes whose copy number ratio are close to the diagonal at values higher or lower than 1, indicating a shared difference to the reference genome, and (iii) genes with a copy number ratio above or below the diagonal, revealing significant gene CNVs between these two isolates(Figure 4B). Both gene depletion and gene amplification were observed in M3015 compared to FJ2002, including increased gene dosage for a biopterin transporter on chr 10 (identified by LtrCDC216_10.0385 and LtrCDC216_10.0390), a histone H4 on chr 21 (identified by LtrCDC216_21.0015), and a hypothetical protein on chr 31 (identified by LtrCDC216_31.1910), or decreased gene dosage for a Isy1-like splicing family on chr 7 (identified by LtrCDC216_07.0020), a tuzin gene on chr 8 (identified by LtrCDC216_08.0795), and a hypothetical protein on chr 13 (identified by LtrCDC216_13.0460) were observed to be depleted in M3015 compared to FJ2002. The biological importance of some of these genes in growth, survival, and gene expression regulation suggests potentially important phenotypic differences between these two isolates.

**Figure 4.**
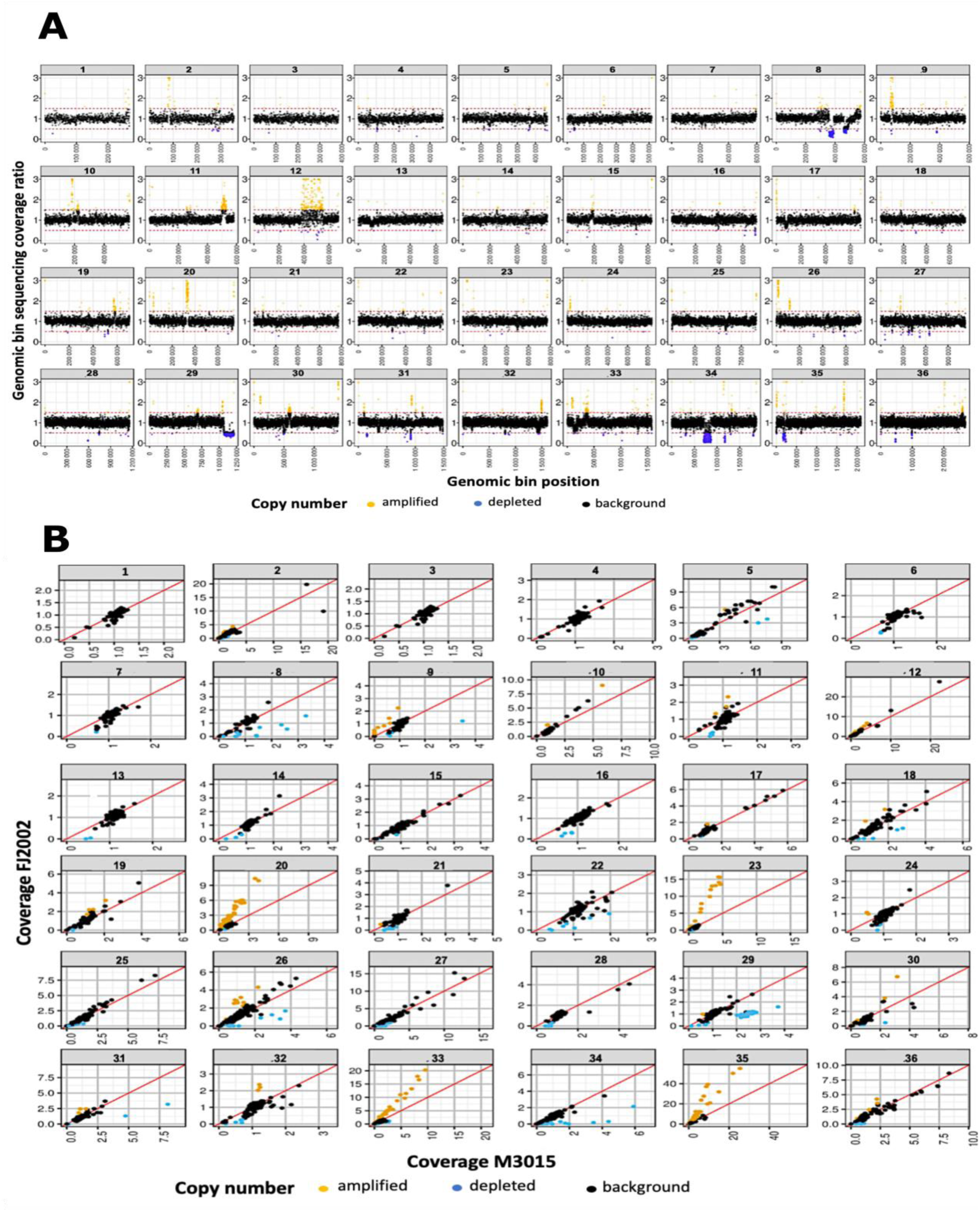
Copy number variation (CNV) analysis. (A) Scatter plots of the normalized sequencing coverage ratios of genomic bins between the FJ2005 and M3015 samples. The X-and Y-axes show respectively coverage ratio and genomic position. Ratio scores >1.25 and <0.75 are indicated in respectively orange and blue. (B) Chromosome-specific scatter plots of gene CNVs between FJ2002 and M3015 isolates. Individual genes are represented by dots. The red diagonal lines indicate the bisectors.

### Comparative SNP analysis in the Moroccan L. tropica isolates

We assessed the diversity of the Moroccan *L. tropica* isolates by analyzing the number, frequency and localization of SNPs. As indicated in Table S4, 178,469 SNPs were identified in Ltr_16 in comparison to the reference genome, while between 98,729 and 111,393 SNPs were identified in the other 13 isolates. We then analyzed unique and shared SNPs between and within the isolates (Figure 5A), revealing high genomic similarity between FJ2005, M3015 and M2013 that share 79,096 SNPs (75%), while only 42% were shared with Ltr_16. Furthermore, Ltr_16 shared 5366 (3%) SNPs with M3015, indicating weak genomic relationship and confirming the evolutionary distance from the other isolates observed by PCA and clustering analyses (see Figure 2).

**Figure 5.**
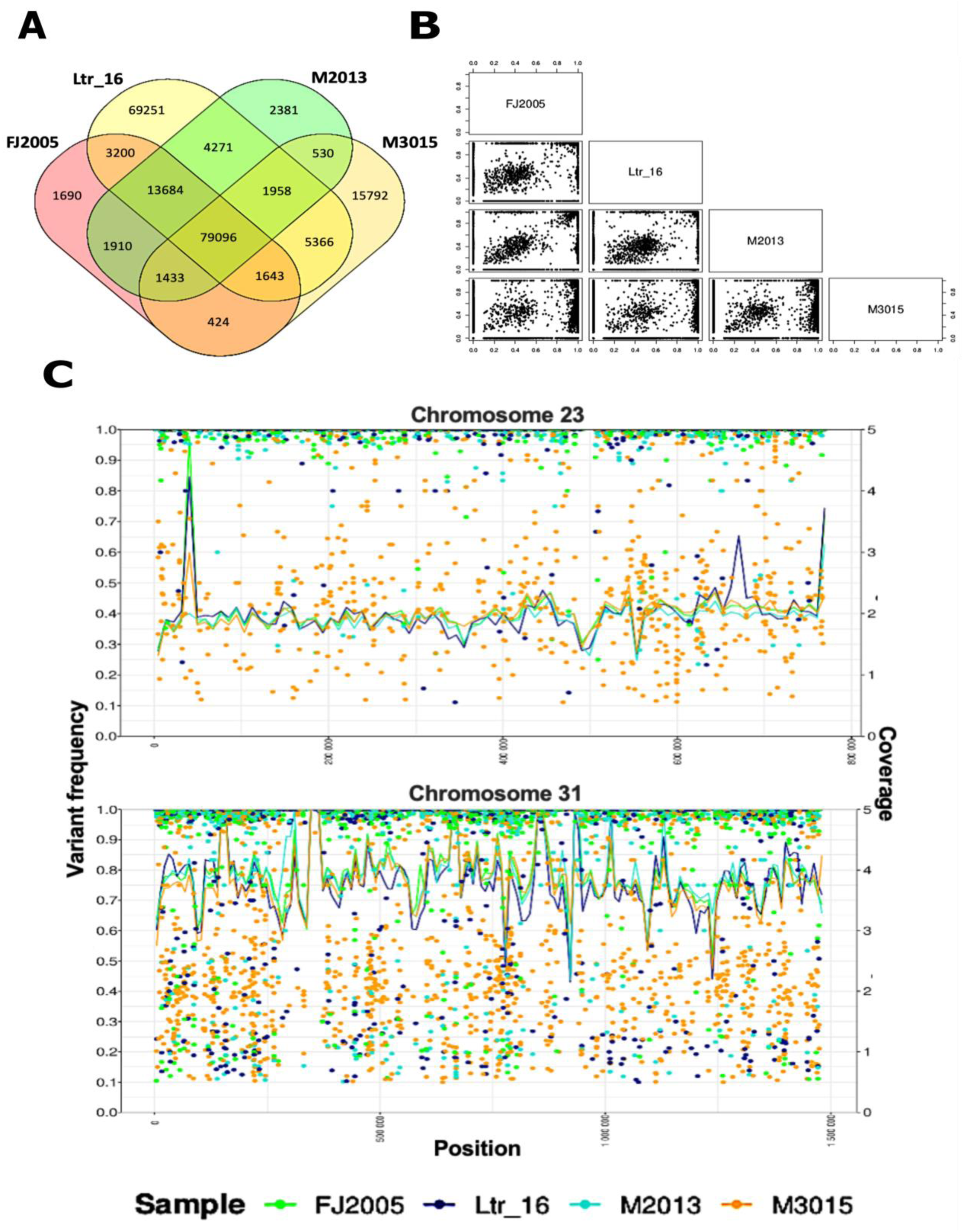
SNP analysis. (A) Venn diagrams showing the number of unique and shared SNPs among four isolates that were isolated in 2020 (FJ2005), 2016 (Ltr_16), 1990 (M2013), and 1995 (M3015). (B) Pairwise scatter plot comparing the variant allele frequency of all SNPs detected in the four isolates FJ2005, Ltr_16, M2013 and M3015. (C) Combined plots of SNP frequency (left Y-axis) and genomic localization (X-axis) (dots), and bin coverage across (right Y-axis, line) of FJ2005, Ltr_16, M2013, and M3015 for chromosomes 23 and 31 is shown.

Direct, pairwise comparison of SNPs frequencies of the four isolates FJ2005, Ltr_16, M2013 and M3015 allowed us visualized the distribution of SNPs. The scatter plots reveal a prominent signal at the upper right corner, indicating shared SNPs at frequencies close to 100% (Figure 5B), as well as horizontal and vertical signals emerging from this signal, representing shared SNPs that are 100% in only one of the two isolates. A second set of shared SNPs was scattered at the center of the plot, indicating SNPs at lower frequencies in both isolates, suggesting the presence of subpopulations that are defined by variations in SNP frequency. Unique SNPs are identified along the vertical and horizontal axes at a frequency of 0 for one of the two isolates. Analyzing the SNP density distribution uncovered a significant number of heterozygous SNPs in isolate M3015 that show a patchy distribution (Supplementary Figure S1 and Figure 5C), which is compatible with a recent hybridization event followed by genetic recombination. Most of the SNPs showed high frequency between 90-100%, which either may be caused by divergent evolution with respect to the reference isolate, or reflect an ancient hybridization event with gene conversion causing fixation (Figure 5C).

### Cluster analysis of the Moroccan L. tropica isolates

We next performed phylogenomic analysis to gain first insight into the possible origin of the divergent isolates Ltr_16 and M3015. Comparing the genomes of our 14 Moroccan isolates with 8 publicly available *L. tropica* genomes from Lebanon, Jordan, and Saudi Arabia (NCBI BioProjects: PRJNA438080, PRJNA453461, and PRJNA589179) again resulted in clustering of the 12 isolates representing the core isolates from the Azilal province. In contrast, Ltr_16 clustered closer to two Lebanese isolates, while M3015 groups closest to Jordanian and Israeli isolates (Figure 6). The large divergence observed between all isolates (except within the core group) suggests that the *L. tropica* genome seems to evolve in a local fashion in response to ecological constraints that still remain to be elucidated.

**Figure 6.**
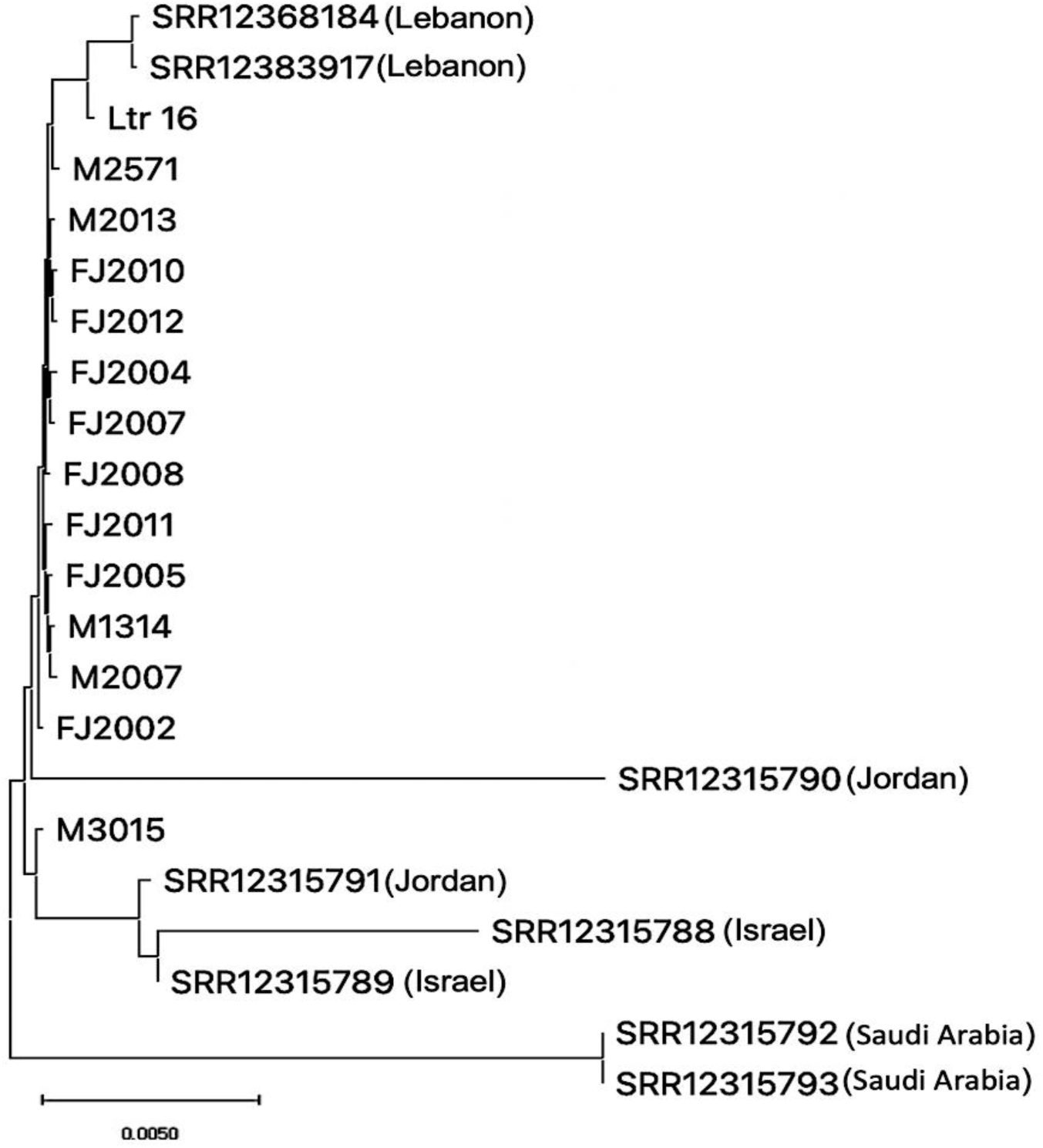
Phylogenomic tree analysis comprising the fourteen Moroccan isolates and an additional eight *L. tropica* genomes from the regions indicated.

## DISCUSSION

We present the first comprehensive genome diversity study of Moroccan *L. tropica* isolates by analyzing high resolution WGS data. SNP analysis of 14 *L. tropica* isolates revealed a genetically highly related core group of 12 isolates that showed significant divergence to the *L. tropica* reference genome generated from a Pakistani isolate [21], and a series of publicly available *L. tropica* genomes from various countries across the Middle-East. Despite the small sample size, this divergence could suggest the existence of local SNP signatures in *L. tropica*, which seem genetically complex in Morocco given the presence of shared SNPs at intermediate and low frequency. We further identified two isolates (Ltr_16 and M3015) that diverged both from the reference genome as well as from the Azilal core group, thus confirming the genetic heterogeneity in-between *L. tropica* isolates in Morocco.

*Leishmania* parasites survived over many millions of years under selective pressures caused by natural ecological changes (storms, floods, hurricanes, volcanic eruptions, etc.), leading to frequent disruption of established host–vector relationships, which may explain today’s genotypic complexity of *Leishmania* vectors and reservoirs. Previous climatic changes, ice ages and the formation of arid regions, as well as land mass disruptions are the likely cause of the *Leishmania* genetic variability we observed today at the species level, which was shaped by genetic drift, natural selection and divergent evolution. Little is known on how *Leishmania* responds to more local changes in ecology through short-term evolutionary processes – an open question of increasing importance given the accelerating ecological changes due to deforestation, global warming or armed conflicts [25]. Adaptation of *Leishmania* to these changes through its intrinsic genome instability may likely have unpredictable consequences on the clinical outcome and the epidemiology of the disease. The genomic divergency we revealed in our study proposes *L. tropica* as an interesting ecological model to study *Leishmania* short-term adaptation under field settings in future investigations.

In Morocco, the epidemiology of CL due to *L. tropica* is not yet fully understood. Although the epidemics caused by these parasites are less severe than those caused by *L. major*, *L. tropica* is perceived as a more important public health threat due to their wide geographic distribution [8, 26, 27]. The genomic diversity we document in our study reveals in addition may point to a higher capacity of *L. tropica* to evolve, which would reinforce the clinical threat posed by this *Leishmania* species. Our study resonates with previous investigations on the genetic diversity and heterogeneity of *L. tropica* isolates in Morocco. First, a study by Krayter et al. (2014) used Multilocus Enzyme Electrophoresis (MLEE) to compare the microsatellite profiles of nine *L. tropica* strains isolated from human cases of CL in two different provinces of Morocco, which were further compared to 147 strains isolated from different geographical locations worldwide. The study found that *L. tropica* strains in each of the two Moroccan provinces separated into two phylogenetic clusters independent of their geographical origin, indicating a high degree of genetic heterogeneity, which was linked to the importation of pre-existing variants of *L. tropica* into Morocco rather than local, divergent evolution [28]. Second, El Hamouchi et al. (2019) analyzed genetic polymorphisms of *L. tropica* strains isolated from 125 cutaneous leishmaniasis patients, which showed a significant correlation between intraspecific *L. tropica* variants and geographic origins of the isolates, and revealed 13 distinct haplotypes thus confirming the genetic heterogeneity of *L. tropica* in Morocco [29]. Finally, a study by EL Kacem et al. (2021) applied multilocus sequence typing (MLST) on 48 samples from CL patients from two different localities in Morocco, again indicating high genetic divergence between and among populations[11].

In conclusion, our study provides first genomic evidence for *L. tropica* genetic heterogeneity within and between endemic regions, which is likely the combined result of genetic drift and geographic adaptation, the latter driven by specific ecological factors related to transmission. Our results open important questions on the evolutionary forces and ecological constraints that drive region-specific adaptation of *Leishmania* in general and *L. tropica* in particular. Future studies applying our comparative genomics approach to a large number of geographically distinct *L. tropica* isolates will need to integrate eco-epidemiological information on sand fly species and host reservoirs to identify gene-environment interactions that may influence the clinical outcome of the disease in a given region and the propensity of the parasite to develop drug-resistant phenotypes.

## ACKNOWLEDGEMENTS

The authors would like to express their gratitude to Dr Jean-Claude Dujardin for taking the time to read our paper and providing valuable constructive criticism. We also wish to thank “the Parasitology, Mycology Department, CHU de Montpellier” for providing us with some *L. tropica* isolates. Additionally, we extend our deep appreciation to the local team of Foum-Jamaa’s Health Centers and the Health Delegation of the Moroccan Ministry of Health for their assistance.

## SUPPLEMENTARY FIGURES LEGEND

**Figure S1:** SNP frequency density plots for M3015, Ltr_16, FJ2005, and M2013. The x-axis reports the variant allele frequency. The y-axis the estimated kernel density between 0 and 4 **(Figure S1.pdf)**.

## SUPPLEMENTARY TABLES LEGEND

**Table S1:** Bin copy number variation **(TableS1.xlsx)**.

**Table S2:** Gene copy number variation **(TableS2.xlsx)**. **Table S3:** Total SNPs in each sample **(TableS3.xlsx)**.

**Table S4:** SNPs frequency in M3015, Ltr_16, FJ2005, and M2013 samples (TableS4.xlsx).

## REFERENCES

1. Gijón-Robles P, Abattouy N, Merino-Espinosa G, El Khalfaoui N, Morillas-Márquez F, et al. Risk factors for the expansion of cutaneous leishmaniasis by Leishmania tropica: Possible implications for control programmes. Transbound Emerg Dis 2018;65:1615–1626.

2. Dereure J, Rioux J-A, Khiami A, Pratlong F, Périères J, et al. Écoépidémiologie des Leishmanioses en Syrie. 2 — Présence, chez le chien, de Leishmania infantum Nicolle et Leishmania tropica (Wright) (Kinetoplastida — Trypanosomatidae). Ann Parasitol Hum Comp 1991;66:252–255.

3. WHO. World Health Organization (WHO). https://www.who.int/news-room/fact-sheets/detail/leishmaniasis (2022, accessed 28 January 2023).

4. Marty P, Le Fichoux Y, Pratlong F, Rioux JA, Rostain G, et al. Cutaneous leishmaniasis due to Leishmania tropica in a young Moroccan child observed in Nice, France. Transactions of the Royal Society of Tropical Medicine and Hygiene 1989;83:510.

5. Pratlong F, Rioux J-A, Dereure J, Mahjour J, Gallego M, et al. Leishmania tropica au Maroc. IV — Diversité isozymique intrafocale. Ann Parasitol Hum Comp 1991;66:100–104.

6. Ait Kbaich M, Mhaidi I, Ezzahidi A, Dersi N, Hamouchi A, et al. New Epidemiological pattern of cutaneous leishmaniasis in two Pre-Saharan arid provinces, southern Morocco. Acta Tropica;173. Epub ahead of print 1 May 2017. DOI: 10.1016/j.actatropica.2017.05.016.

7. El Idrissi Saik I, Benlabsir C, Fellah H, Lemrani M, Riyad M. Transmission patterns of Leishmania tropica around the Mediterranean basin: Could Morocco be impacted by a zoonotic spillover? PLoS Negl Trop Dis 2022;16:e0010009.

8. Guessous-Idrissi N, Chiheb S, Hamdani A, Riyad M, Bichichi M, et al. Cutaneous leishmaniasis: an emerging epidemic focus of Leishmania tropica in north Morocco. Transactions of the Royal Society of Tropical Medicine and Hygiene 1997;91:660–663.

9. Lemrani M, Nejjar R, Pratlong F. A new Leishmania tropica zymodeme--causative agent of canine visceral leishmaniasis in northern Morocco. Ann Trop Med Parasitol 2002;96:637– 638.

10. El Hamouchi A, El Kacem S, Ejghal R, Lemrani M. Genetic polymorphism in Leishmania infantum isolates from human and animals determined by nagt PCR-RFLP. Infect Dis Poverty 2018;7:54.

11. El Kacem S, Kbaich MA, Daoui O, Charoute H, Mhaidi I, et al. Multilocus sequence analysis provides new insight into population structure and genetic diversity of Leishmania tropica in Morocco. Infect Genet Evol 2021;93:104932.

12. Metzker ML. Sequencing technologies - the next generation. Nat Rev Genet 2010;11:31– 46.

13. Bussotti G, Gouzelou E, Boité MC, Kherachi I, Harrat Z, et al. Leishmania Genome Dynamics during Environmental Adaptation Reveal Strain-Specific Differences in Gene Copy Number Variation, Karyotype Instability, and Telomeric Amplification. mBio;9. Epub ahead of print 21 December 2018. DOI: 10.1128/mBio.01399-18.

14. Bussotti G, Benkahla A, Jeddi F, Souiaï O, Aoun K, et al. Nuclear and mitochondrial genome sequencing of North-African Leishmania infantum isolates from cured and relapsed visceral leishmaniasis patients reveals variations correlating with geography and phenotype. Microb Genom;6. Epub ahead of print October 2020. DOI: 10.1099/mgen.0.000444.

15. Bussotti G, Piel L, Pescher P, Domagalska MA, Rajan KS, et al. Genome instability drives epistatic adaptation in the human pathogen Leishmania. Proceedings of the National Academy of Sciences 2021;118:e2113744118.

16. Prieto Barja P, Pescher P, Bussotti G, Dumetz F, Imamura H, et al. Haplotype selection as an adaptive mechanism in the protozoan pathogen Leishmania donovani. Nat Ecol Evol 2017;1:1961–1969.

17. Schwabl P, Boité MC, Bussotti G, Jacobs A, Andersson B, et al. Colonization and genetic diversification processes of Leishmania infantum in the Americas. *Commun Biol* 2021;4:139.

18. Downing T, Imamura H, Decuypere S, Clark TG, Coombs GH, et al. Whole genome sequencing of multiple Leishmania donovani clinical isolates provides insights into population structure and mechanisms of drug resistance. Genome Res 2011;21:2143–2156.

19. Dumetz F, Cuypers B, Imamura H, Zander D, D’Haenens E, et al. Molecular Preadaptation to Antimony Resistance in Leishmania donovani on the Indian Subcontinent. mSphere 2018;3:e00548-17.

20. Späth GF, Bussotti G. GIP: An open-source computational pipeline for mapping genomic instability from protists to cancer cells. 2021;2021.06.15.448580.

21. Unoarumhi Y, Batra D, Sheth M, Narayanan V, Lin W, et al. Chromosome-Level Genome Sequence of Leishmania (Leishmania) tropica Strain CDC216-162, Isolated from an Afghanistan Clinical Case. Microbiol Resour Announc 2021;10:e00842–20.

22. Steinbiss S, Silva-Franco F, Brunk B, Foth B, Hertz-Fowler C, et al. Companion: a web server for annotation and analysis of parasite genomes. Nucleic Acids Res 2016;44:W29– W34.

23. Iantorno SA, Durrant C, Khan A, Sanders MJ, Beverley SM, et al. Gene Expression in Leishmania Is Regulated Predominantly by Gene Dosage. mBio 2017;8:e01393–17.

24. Rogers MB, Hilley JD, Dickens NJ, Wilkes J, Bates PA, et al. Chromosome and gene copy number variation allow major structural change between species and strains of Leishmania. Genome Res 2011;21:2129–2142.

25. Shaw JJ, De Faria DL, Basano SA, Corbett CE, Rodrigues CJ, et al. The aetiological agents of American cutaneous leishmaniasis in the municipality of Monte Negro, Rondônia state, western Amazonia, Brazil. Ann Trop Med Parasitol 2007;101:681–688.

26. Guernaoui S, Boumezzough A, Laamrani A. Altitudinal structuring of sand flies (Diptera: Psychodidae) in the High-Atlas mountains (Morocco) and its relation to the risk of leishmaniasis transmission. Acta Trop 2006;97:346–351.

27. Guilvard E, Rioux JA, Gallego M, Pratlong F, Mahjour J, et al. [Leishmania tropica in Morocco. III--The vector of Phlebotomus sergenti. Apropos of 89 isolates]. Ann Parasitol Hum Comp 1991;66:96–99.

28. Krayter L, Alam MZ, Rhajaoui M, Schnur LF, Schönian G. Multilocus Microsatellite Typing reveals intra-focal genetic diversity among strains of Leishmania tropica in Chichaoua Province, Morocco. Infection, Genetics and Evolution 2014;28:233–239.

29. El Hamouchi A, Ajaoud M, Arroub H, Charrel R, Lemrani M. Genetic diversity of Leishmania tropica in Morocco: does the dominance of one haplotype signify its fitness in both predominantly anthropophilic Phlebotomus sergenti and human beings? Transboundary and Emerging Diseases 2019;66:373–380.

