## Supplemental Figure S1 for "A comparative genomics approach reveals a local genetic signature of *Leishmania tropica* in Morocco"

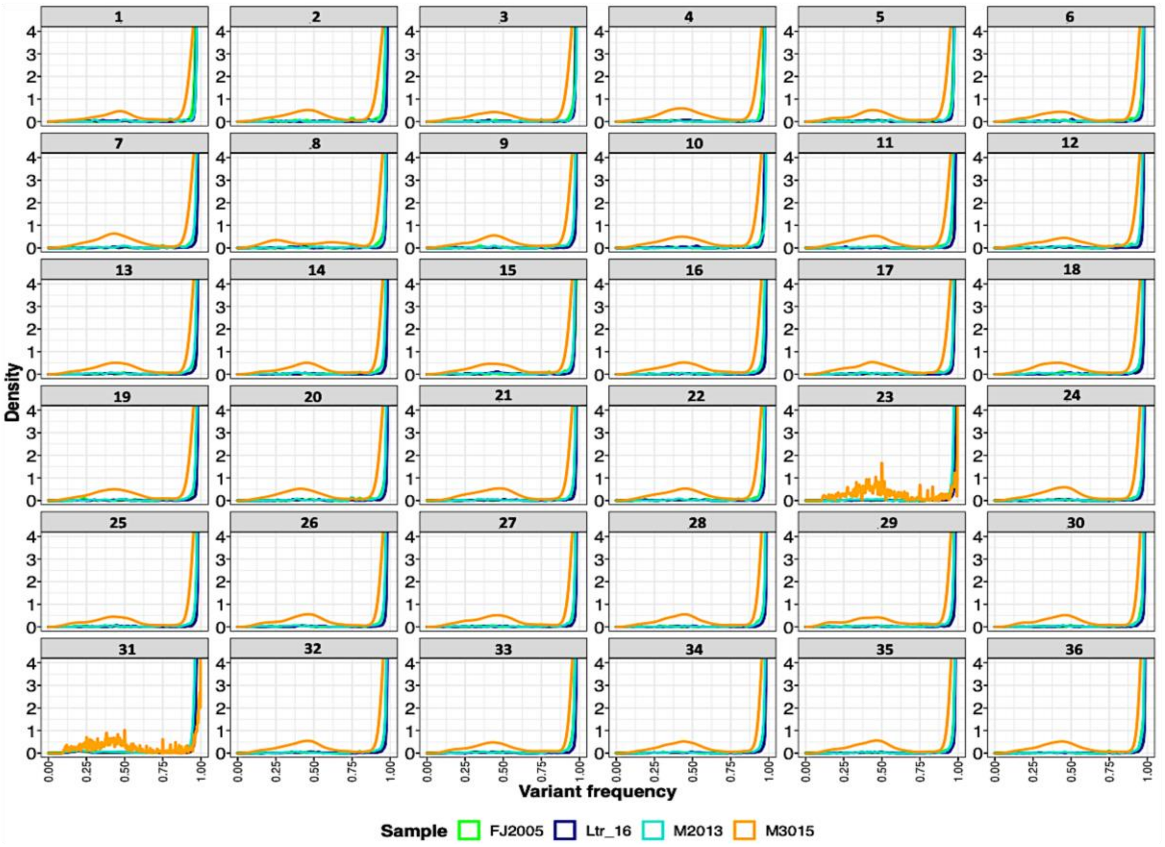


*Figure S1. SNP frequency density plots for M3015, Ltr_16, FJ2005, and M2013. The x-axis reports the variant allele frequency. The y-axis the estimated kernel density between 0 and 4.*
